# A causal study of bumetanide on a marker of excitatory-inhibitory balance in the human brain

**DOI:** 10.1101/2020.09.22.304279

**Authors:** Thomas L. Botch, Alina Spiegel, Catherine Ricciardi, Caroline E. Robertson

## Abstract

Bumetanide has received much interest as a potential pharmacological modulator of the putative imbalance in excitatory/inhibitory (E/I) signaling that is thought to characterize autism spectrum conditions. Yet, currently, no studies of bumetanide efficacy have used an outcome measure that is modeled to depend on E/I balance in the brain. In this manuscript, we present the first causal study of the effect of bumetanide on an objective marker of E/I balance in the brain, binocular rivalry, which we have previously shown to be sensitive to pharmacological manipulation of GABA. Using a within-subjects placebo-control crossover design study, we show that, contrary to expectation, acute administration of bumetanide does not alter binocular rivalry dynamics in neurotypical adult individuals. Neither changes in response times nor response criteria can account for these results. These results raise important questions about the efficacy of acute bumetanide administration for altering E/I balance in the human brain, and highlight the importance of studies using objective markers of the underlying neural processes that drugs hope to target.

## Introduction

Excitatory and inhibitory (E/I) activity is balanced in neural systems at multiple spatial scales [1, 2], and this balance is thought to be critical for typical neural function [3–5]. Multiple lines of evidence implicate disrupted E/I balance in the neurobiology of Autism Spectrum Conditions (ASC; autism henceforth) [6–12]. In particular, studies in both humans and in animal models suggest that altered inhibitory signaling, mediated by the neurotransmitter GABA, may characterize the condition [10, 11]. Despite the accumulating evidence, the intricacies of autism neurobiology are poorly understood, hindering efforts to develop treatment strategies for the condition.

One prominent developmental account of autism proposes a disruption of an important neurobiological milestone, known as the GABA-switch, as a potential explanation for disturbed inhibitory action in the autistic brain [13]. During development, the polarity of GABAergic action transitions from excitatory (depolarizing) to inhibitory (hyperpolarizing) due to a progressive reduction in intracellular chloride (Cl-) concentration in principal neurons [14, 15] -- a developmental sequence that may be disrupted in animal models of autism [16, 17]. In light of these accounts, it has been posited that augmenting GABAergic action might provide a promising therapeutic for some symptoms associated with autism [13, 18].

Bumetanide, a loop diuretic, has proven hopeful in rectifying GABA polarity in valproic acid and Fragile X animal models of autism [16, 19]. Bumetanide is thought to increase the hyperpolarizing potential of GABA by blocking NKCC1 receptors, which are responsible for Cl-entrance into the cell [20]. Further, some studies of bumetanide in humans, specifically children with autism, have shown evidence for attenuation of social symptom severity and improvement of emotion recognition [21–23], although, notably, these benefits are not universally observed [24]. Importantly, to date, direct evidence that bumetanide increases inhibition in the human brain is lacking, which complicates linking the reported symptomatic benefits to the drug’s predicted physiological effects.

Therefore, we sought to test the effects of bumetanide on a robust behavioral index of E/I balance, binocular rivalry. Rivalry is a simple visual phenomenon that is modeled to rely on the on the balance of inhibition and excitation in visual cortex [25–30]. Prior pharmacological studies in humans reveal a causal link between rivalry dynamics and GABAergic inhibition using both GABA_A_ and GABA_B_ modulators [31, 32], as well as a dependence of rivalry dynamics on tonic levels of GABA in visual cortex [11, 32]. Given these links between rivalry dynamics and E/I balance in visual cortex, as well as recent evidence showing altered rivalry dynamics in adult individuals with autism [11, 33–35], rivalry has been suggested as a noninvasive perceptual marker of E/I signaling in visual cortex, and its putative disturbance in psychiatric conditions, including autism.

Here, we asked whether acute bumetanide administration would alter rivalry dynamics. We hypothesized that bumetanide would increase the degree to which individuals predominantly perceive one image fully suppressed from awareness (“perceptual suppression”), which computational and empirical data suggest is gated by GABAergic inhibition [31, 36, 37]. We tested this hypothesis in a within-subjects drug-placebo, cross-over design pharmacological study of rivalry dynamics in neurotypical adults.

## Materials and Methods

### Participants

21 healthy adults (N = 15 female; mean age 22.5 +/-3.68 SD years) participated in the study. Written consent was obtained from all participants, and all studies were approved by the Massachusetts Institute of Technology Institutional Review Board. All participants had normal or corrected-to-normal vision, were neither pregnant nor nursing, and were free from: (1) any known history of psychiatric or neurological conditions; (2) any other diagnosed medical conditions, including a history of heart failure; (3) any psychiatric medications; and (4) any known drug allergies (including bumetanide). All studies took place at the MIT Clinical Research Center, under the constant observation of a research nurse/nurse practitioner (C.R.) and nursing team.

### Study drugs: bumetanide (loop-diuretic)

Participants participated in a study investigating the effects of bumetanide (1 mg) on binocular rivalry dynamics. Bumetanide is an FDA-approved loop-diuretic known to antagonize sodium-potassium-chloride cotransporters, NKCC1 and NKCC2, which modulate intracellular chloride concentration. At low concentrations, bumetanide has a high affinity to block NKCC1, thereby reducing intracellular chloride concentration and, by proxy, altering GABAergic action potentials [14, 20]. Bumetanide dosage was chosen to fall within the standard prescribed range.

### Experimental design: placebo-controlled crossover design

Each participant took part in a 3-day study, comprised of: a health assessment/practice session (Day 1) and 2 experimental days (Days 2 and 3). On each experimental day, a participant was given either a drug or a placebo and participated in a short binocular rivalry experiment after the drug had come into effect (Fig. 1*B*). This within-participant crossover design allowed us to compare the effects of each drug, relative to those of a placebo, on rivalry dynamics. The design of this study was identical to that used in a recent study of the effects of GABA_A_ and GABA_B_ modulators on rivalry dynamics [31].

**Figure 1.**
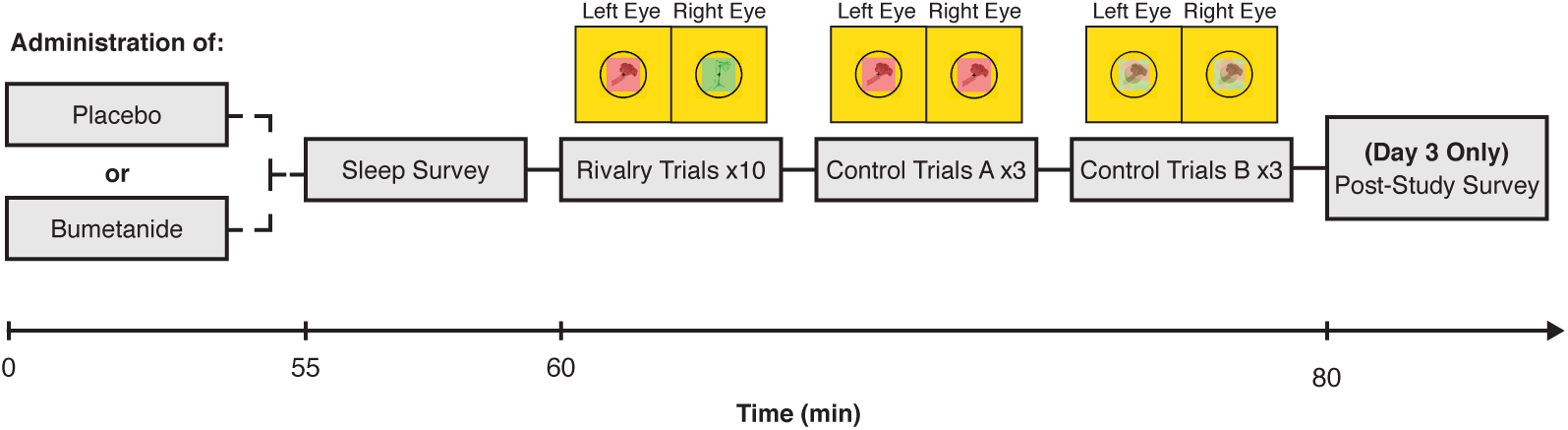
Experimental paradigm. Participants took part in a double-blind crossover study to test the effects of bumetanide, a drug thought to alter GABAergic action, on binocular rivalry, a noninvasive marker of excitatory-inhibitory balance in visual cortex. After an initial study visit (Day 1; see Materials and Methods), participants returned for two experimental testing days (Day 2 and Day 3). On each experimental day, participants were administered either a placebo or the drug. After waiting for drug absorption, participants completed binocular rivalry and control rivalry replay trials, so that the effect of bumetanide on rivalry dynamics could be assessed.

On Day 1, participants were briefed on the study and consented to participate. Next, the nurse practitioner (NP) assessed the participant for drug allergies, reviewed the participant’s medical history, assessed the participant for colorblindness, and completed a routine physical assessment (including blood pressure, temperature, and a basic metabolic panel to assess kidney function) to ensure that participants were healthy enough to participate in the study. All women of childbearing ages were asked to provide a urine specimen for a pregnancy point of care test to confirm non-pregnancy state. Finally, participants performed six 45 s rivalry practice trials to become familiarized with the task.

On Days 2 and 3 of testing, each participant received a pill: an oral dose of a drug on one day and a matched placebo pill on the other day (Figure 1). Drug day assignment was blinded, randomized, and counterbalanced across participants (i.e., half of the participants received a placebo on Day 2 and bumetanide on Day 3, whereas the other half received bumetanide on Day 2 and a placebo on Day 3). After waiting for a time period to allow for drug absorption and distribution to approximate peak-plasma concentration (60 min; [38]), the participants completed a brief drowsiness scale and performed the binocular rivalry experiment. After testing was complete on each day, the NP performed a physical assessment for adverse events to ensure that each participant was in sufficient health to leave the Clinical Research Center. Additionally, participants answered a series of survey questions regarding their first-person reflections on their rivalry dynamics during the study at the end of Day 3.

### Experimental paradigm

Procedures and stimuli for the binocular rivalry experiments matched those used in a recent study of the effects of GABA_A_ and GABA_B_ modulators on rivalry dynamics [31] (see reference for more details). In brief, each experimental session consisted of two blocks of five 45 s experimental runs (binocular rivalry), and six 45 s rivalry replay control trials (3 “sudden onset” trials and 3 “gradual onset” trials). Note, due to time constraints only a subset of participants completed the rivalry replay trials. During rivalry trials, participants viewed two different images (red or green), one image presented to each of the participant’s eyes. During rivalry replay trials, participants viewed two identical images which were continuously presented to both eyes and alternated temporally on the screen to mimic perceptual shifting during rivalry. In all trials. participants were asked to continuously report whether they perceived a fully dominant percept, the red image (right key) or the green image (left key), or a mixture of the two images (up key). Percept durations as well as switch rates were analyzed.

### Binocular rivalry experiment: experiment and analysis

As in [31], on each trial, two grayscale objects (e.g., a baseball and a piece of broccoli) appeared on the left and right of the screen. Each object was displayed within a tinted square (green or red; width: 5.3°), surrounded by a black circle to support binocular fusion (radius: 3.7°). Participants were instructed to fixate on a cross at the center of each image. Stimuli were viewed through a head-mounted display (Oculus Rift CV1; 1080 × 1200 pixels per eye, 90 Hz refresh rate, FOV = 110°), which displayed the left and right halves of the screen to the participants’ left and right eyes such that each eye was presented with only one of the two images (red or green; Fig. 1*B*). Responses were collected via a keyboard. A sequence of perceptual events was later computed based on when one continuous key press was terminated and another began. Keypresses lasting <450 ms or periods where no key was pressed were omitted from the analysis. For each participant and trial, the frequency of perceptual transitions as well as the duration of any perceptual event (red, green, or mixed) were calculated. Transitions were subdivided into “switches” (e.g., red to mixed to green) and “reversions” (e.g., red to mixed to red).

### Rivalry replay trials: paradigm and analysis

Binocular rivalry replay trial stimuli were identical to those used in the main rivalry experiment, and the paradigm was identical to our previously published studies [31, 33]. In brief, participants viewed a simulated series of back-and-forth binocular rivalry alternations. These trials allowed us to measure two non-perceptual factors which could potentially contribute to any observed drug effects on rivalry: response latencies (Control A) and response criteria (Control B). In Control A trials (“sudden-onset”), participants viewed stepwise, sudden transitions between two stimuli, which allowed us to determine response latencies to clear, obvious transitions. In Control B trials (“gradual-onset”), participants viewed gradual, linear transitions between stimuli created using a series of dynamic Gaussian image filters. This allowed us to determine participants’ response criteria: the extent of image blending each individual required to identify an image as mixed versus dominant. See [31] for more details.

### Perceptual suppression metric

Perceptual suppression during rivalry is experienced by the viewer as the percept of a single image (while another is fully suppressed from awareness). Thus, the proportion of perceptual suppression, was calculated as the proportion of each trial spent viewing a fully dominant percept: (dominant percept durations)/(dominant + mixed percept durations).

### Statistical analyses

Two-tailed, uncorrected *p* values and estimates of effect sizes (*r*) are reported for all effects. In all rivalry analyses, Wilcoxon signed-rank tests were used to compare drug vs. placebo effects, as the distribution rivalry percept durations were not normally distributed (Shapiro-Wilk test; *p* < 0.05). Spearman’s rank correlation coefficients (*R*s) are reported for reliability. For each participant, trials where rivalry was never initiated (one percept was reported for the entire trial) were eliminated. Participants missing >50% of trials (N=0) or whose rivalry percept durations were determined to fall outside of 2 SD of the group mean (N=1) were excluded from all analyses.

### Karolinska Sleepiness Scale

Immediately before each rivalry session, participants completed the Karolinska sleepiness scale (KSS) to assess any self-reported changes in drowsiness between drug and placebo testing sessions. The KSS consisted of a single question, “Choose the position on the following scale that best describes your current state”, where answers ranged from 1 (extremely alert) to 9 (very sleepy, great effort to keep awake, fighting sleep).

### Experience questionnaire

After the final testing session on Day 3, participants completed a post-study survey to assess: (1) “On which day do you think you received the drug and why?”, (2) “Did you notice any differences in your experience of binocular rivalry between yesterday and today? If so, please describe.”, (3) “How confident were you in determining whether you were seeing a mixture or a dominant state?”, and (4) “Describe what mixtures during binocular rivalry look like to you.”

## Results

We predicted that bumetanide, a drug known to alter intracellular Cl-concentration and, by proxy, posited to increase GABAergic inhibition, would increase perceptual suppression during rivalry. We also assessed performance on rivalry replay control trials to establish whether any observed changes were due to non-perceptual effects on response latencies or response criteria [39, 40].

### Bumetanide does not alter perceptual suppression

To test whether bumetanide affects the depth of perceptual suppression during rivalry, we calculated the drug effect on the proportion of suppression for each individual (Proportion of Suppression on Drug - Placebo days) using a Wilcoxon signed-rank test. We observed a trend towards lower proportions of perceptual suppression on the drug, compared to the placebo day (Z = −1.643, *p* = 0.10, *r* = 0.367; Fig 2a). This trend was driven by decreased time in dominant percepts on the drug day compared to the placebo day (Z = −2.315, *p* = 0.021, *r* = 0.518; Fig 2a, top); no change to mixed percepts was observed (Z = −0.597, *p* = 0.55, *r* = 0.133; Fig 2a, bottom). The absence of a drug effect on perceptual suppression was stable across the experimental testing session: perceptual suppression did not significantly differ between the first half (trials 1-5) and second half (trials 6-10) of trials in the experiment (Z = −1.195, *p* = 0.232, *r* = 0.267; Fig 2b). These results indicate that bumetanide does not influence a marker of inhibitory competitive dynamics in visual cortex, perceptual suppression. This finding stands in contrast to studies showing that GABA_A_ and GABA_B_ modulators increase perceptual suppression during rivalry [31].

**Figure 2.**
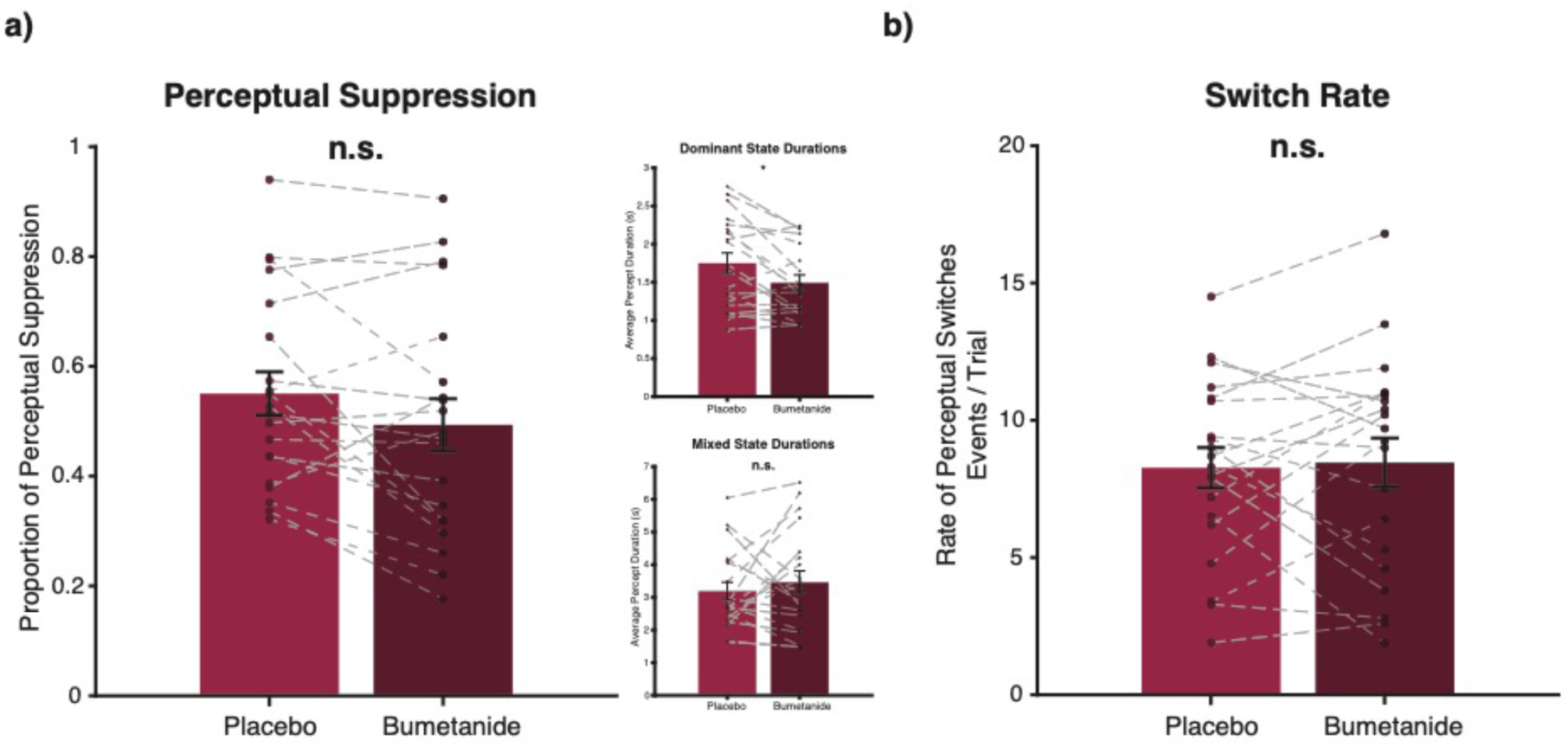
Experimental results. **a)** No effect of bumetanide administration was found on perceptual suppression (Z = −1.634, p = 0.10, *r* = 0.367), although this effect trended in the opposite direction as predicted. (top) This trend was driven by less time in dominant percepts (Z = −2.315, p = 0.021, *r* = 0.518); (bottom) no differences in mixed percept durations were observed (Z = −0.597, p = 0.55, *r* = 1.133). **b)** No effect on switch-rate was observed (Z = −0.28, p = 0.779, *r* = 0.062).

### Bumetanide does not alter rivalry switch rates or reversions

Bumetanide did not reliably affect switch rates (Fig. 2b, Bumetanide: 8.27 +/-0.75 SE switches; Placebo: 8.46 +/-0.91 SE switches, Z = −0.280, *p* = 0.779, *r* = 0.062) or reversion rates (Z = −1.23, *p* = 0.229, *r* = 0.269) during binocular rivalry.

### Drug effects are not confounded by shifts in response latency or response criteria

We next sought to determine whether the drug affected participants’ response latencies or response criteria, as these non-perceptual factors could potentially impact perceptual reports during rivalry. We analyzed performance on two rivalry replay control conditions, which only a subset of took part in due to time constraints: 1) sudden transition trials (N=8), which provided us with an estimate of participants’ motor latencies (how quickly they press a button in response to a stimulus change on the display) and 2) gradual transition trials (N=13), which provided us with an estimate of participants’ perceptual decision criteria (the percentage of mixture in a displayed stimulus for a participant to report either a mixed or dominant percept). A Wilcoxon signed-rank test revealed no effects of drugs on behavior for either response times (Z = −1.54, *p* = 0.123, *r* = 0.544) or decision criteria (Z = −0.314, *p* = 0.753, *r* = 0.087).

### Test-retest reliability

To examine the stability of our primary measure, perceptual suppression, we calculated test-retest reliability by correlating performance on drug versus placebo days across individuals in each study. Participants’ rivalry dynamics were stable across testing days (*R*s = 0.657, *p* = 0.002; Fig. 3). For example, a fast rivaler on Day 2 remained one of the fastest on Day 3. This finding is consistent with previous binocular rivalry studies which illustrate high test-retest reliability of rivalry dynamics within individuals [11, 31, 41].

**Figure 3.**
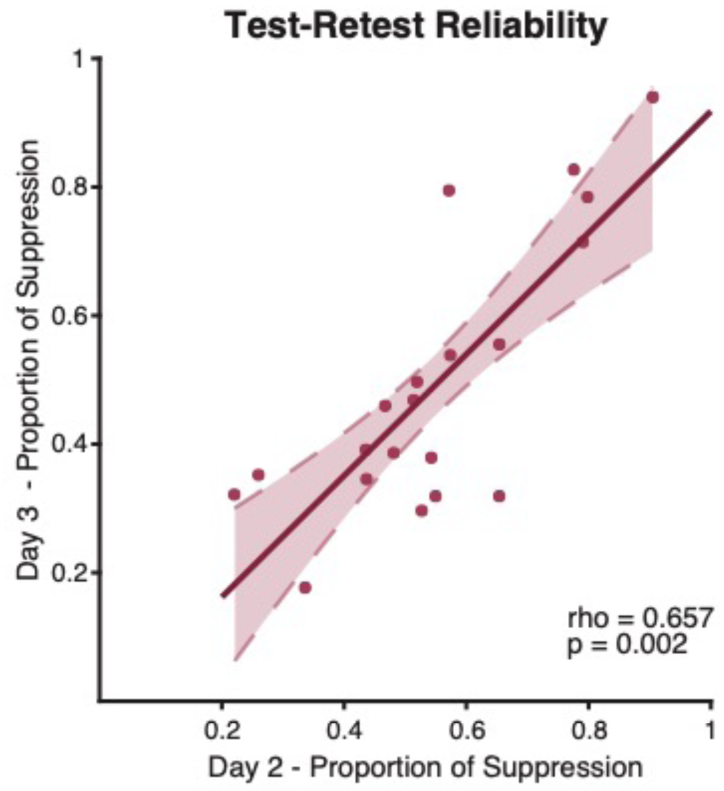
High test-retest reliability of rivalry dynamics. Individual rivalry dynamics were stable across bumetanide and placebo days (Rs = 0.657, p = 0.002). For example, a fast rivaler on Day 2 remained a fast rivaler on Day 3 relative to the group. Dashed lines represent the 95% confidence interval of the regression line.

### Bumetanide does not affect self-reported drowsiness

Participants did not report significant differences in drowsiness between placebo and drug days (mean: 0.35 questionnaire points +/-1.69 points, *p* = 0.367). Therefore, alertness is unlikely to be a contributing factor to these results.

### Participant’s self-reports of drug detection were based on diuretic effects

It is important to consider that participant self-reports, collected via post-study survey, indicated moderate detection of bumetanide administration. Specifically, when asked “on which day do you think you took the drug?”, the majority of participants were correct in their guess: 70%. Critically, most participants based their guess on the diuretic effects (50%), as compared to perceptual changes (20%), cognitive effects (e.g. alertness, sleepiness; 20%), or other reports (e.g. pill color, guessing; 10%).

## Discussion

We have shown that acute administration of bumetanide does not alter binocular rivalry dynamics in neurotypical adult individuals. Indeed, the effects we observed (lower perceptual suppression) here trended in the opposite direction as predicted from previous studies of the impact of GABA modulators on rivalry dynamics [31, 32]. Importantly, bumetanide affected neither of our control measures: neither reaction times to report nor response criteria to determine perceptual transitions in the subset of our participants who participated in rivalry replay trials, thereby ruling out these potentially confounding factors. Our findings raise questions about the efficacy of single-dose administration of bumetanide for altering E/I balance in the human brain.

Recent pharmacological studies confirm that acute modulation of E/I balance using GABAergic modulators alters rivalry dynamics [31, 32]. Two studies from our lab, which used the same rivalry paradigm and placebo-control cross-over pharmacological stutdy design as we used here, found that acute administration of both the GABA_A_ modulator, clobazam, and the GABA_B_ modulator, arbaclofen, increased the strength of perceptual suppression during binocular rivalry relative to placebo [31]. These findings provide causal validation for predictions of computational models of rivalry, which hold that the strength of lateral inhibition between neurons tuned to represent signals from the left vs. right eye percepts governs the strength of perceptual suppression during rivalry [36, 37]. Empirical data using magnetic resonance spectroscopy (MRS) lend further support to these claims: across individuals, the concentration of the neurotransmitters GABA and glutamate(+glutamine) in early visual cortex strongly predict the degree of perceptual suppression [11] and longer dominant percept durations [32, 42] during rivalry. Together, these results suggest that, in principle, binocular rivalry and possibly other related tasks could provide an objective index of GABAergic drug response in the brain. However, our current results question the suitability of rivalry in the case of bumetanide.

Previous studies examining the longitudinal effects of bumetanide in individuals with autism have often demonstrated success in modulating social processing. Specifically, longitudinal bumetanide administration has been observed to improve social domain scores measured by the Child Autism Rating Scale (CARS) [21, 23, 43]. Additionally, recent neuroimaging studies report increased neural response to emotional faces, and normalization of amygdala activation in response to eye-contact in individuals with autism following bumetanide administration [22, 44]. Notably, however, these benefits are not universally observed: improvement of CARS scores is inconsistent across studies [45]. Further, a recent large-scale clinical trial found no improvements in social or sensory symptoms after longitudinal bumetanide treatment in children with autism, as measured using the Social Responsiveness Scale-2 (SRS-2) and Sensory Profile (SP-NL) respectively, and only minor reductions in repetitive behaviors, measured by the Repetitive Behavior Scale Revised (RBS-R) [24]. Taken together, these mixed findings highlight the need to better understand bumetanide’s ability to ameliorate the multiplexed symptoms of autism, and underscore the importance of objective measures of drug efficacy.

It is thought that bumetanide may affect neural processing by modulating E/I balance in the brain. However, our results, along with others in the literature, highlight the need to understand the nuances of bumetanide’s action on E/I balance and cognitive function in humans. One recent MRS study suggests that bumetanide may, as predicted, modulate E/I balance in adults, as evidenced through changes in the GABA-Glutamate ratio in visual and insular cortex pre-relative to post-3-months of bumetanide administration in children with autism [46]. Importantly, however, this study did not report consistent changes in either GABA or Glutamate alone – only in the E/I ratio as indexed by these neurotransmitter levels. Further, the direction of these changes was the opposite of what might be predicted if bumetanide were to increase MRS-based markers of inhibition [13]: bumetanide lowered the E/I ratio, resulting in greater excitation relative to inhibition. Interestingly, our study also observed a non-significant trend towards bumetanide reducing perceptual suppression in control individuals, suggesting that the drug may in fact reduce, rather than increase inhibition in visual cortex. Additional research is needed to unpack the effects of bumetanide on E/I balance in the human brain, using objective markers of drug response in a longitudinal study. Further, it is possible that effects may differ between individuals with and without autism, if the relative intracellular Cl-concentration is indeed affected in autism.

A few other nuances of bumetanide’s mechanism of action are important to consider. First, chronic administration of bumetanide produces substantial diuretic effects and systemic build-up in the kidneys, and may alter neuronal function by way of larger systemic changes [47, 48]. Second, the diuretic effects of bumetanide present a challenge for a truly blinded study: even in our acute administration study, diuresis on the drug day was self-reported by 50% of participants, potentially compromising the blinded nature of this and other studies of bumetanide. Third, although bumetanide demonstrates high specificity as an NKCC1 antagonist [20], NKCC1 is differentially expressed throughout the cortex during development [49, 50], and becomes increasingly present in the PNS throughout maturation [51]. Thus, it is unclear how bumetanide impacts the nervous system, and whether bumetanide exerts differential effects on different areas in the brain. Further characterization of the diverse molecular profile of bumetanide across developmental timepoints is important for understanding its therapeutic potential, and future research should be directed to understand the effects of bumetanide on distinct cortical circuits maintained by E/I balance.

Null results are difficult to interpret; there are multiple reasons why we might not have observed a significant result in our study, as might have been predicted. First, bumetanide exhibits poor transport across the blood-brain barrier (BBB) [52, 53]. Thus, in contrast to other GABA modulators [31], chronic administration may be important to detect an effect of bumetanide on rivalry. Indeed, single-dose administration of bumetanide in animal models have produced low cortical concentrations [52]. Second, while the administered dose of bumetanide is sufficient for peak diuretic properties [38] and comparable to previous longitudinal studies in humans [23], it may fall short of the amount required to exhibit effects on cognitive function.

Looking forward, a major impediment to drug development in psychiatric research is the lack of robust, objective markers of the underlying targeted neural processes [54, 55]. The potential for sensory tasks to provide such markers is particularly high, given their suitability for translational research [56] and presence in conditions such as autism in some [57–60], although not all studies [36, 61]. Given recent links between GABAergic inhibition and rivalry dynamics [11, 31], and the replicated differences in binocular rivalry dynamics observed in individuals with autism [11, 33–35], rivalry has been suggested as a noninvasive perceptual marker of inhibitory signaling in visual cortex, and its putative disturbance in autism [11, 35]. However, further research is needed to establish whether acute bumetanide administration alters such noninvasive markers of E/I balance in humans.

## Acknowledgements

This work was supported by a grant from the Simons Foundation Autism Research Initiative (SFARI #597694) to C.E.R. We thank Nwamaka Amobi and Tatiana Urman for their help with data collection. The authors declare no competing financial interests.

## Author contributions

C.E.R. and C.R. designed research; A.S., C.R., and C.E.R. performed research; C.E.R. and T.L.B. analyzed data; T.L.B. and C.E.R. wrote the paper.

## Citation gender diversity statement

Recent work in several fields of science has iden tified a bias in citation practices such that papers from women and other minorities are under-cited relative to the number of such papers in the field [62–66]. Here we sought to proactively consider choosing references that reflect the diversity of the field in thought, fo rm of contribution, gender and other factors. We obtained predicted gender of the first a nd last author of each reference by using databases that stores the probability of a nam e being carried by a woman [66, 67]. By this measure (and excluding self-citations to the first and last authors of our current paper), our references contained 3.8% woman(first)/ woman(last), 3.8% man/woman, 22.6% woman/man, 69.8% man/man, and 0% unknown categorization. This method is limited in that a) names, pronouns, and social media profiles used to construct the databases may not, in every case, be indicative of gender identity and b) it cannot account intersex, non-binary, or transgender people. We look forw ard to future work that could help us to better understand how to support equitable practi ces in science.

